# Competitive release from native range does not predict invasive success of a globally dominant exotic ant species

**DOI:** 10.1101/239277

**Authors:** Senay Yitbarek, Ivette Perfecto, John H. Vandermeer

## Abstract

A major goal of invasion biology is to understand under what conditions exotic species thrive in the introduced range. High competitive abilities are thought to be an important characteristic of exotic species. Most invasion studies focus on the competitive ability of exotic species in the introduced range and attribute their ecological success to competitive release, but fewer studies have compared the relative competitive differences within the native range. These comparative studies are important in order to determine if competitive abilities of exotic species are strong predictors of invasion success. The little fire ant *Wasmmnia auropunctata* is a highly invasive species that has spread from its original range (Central and South America) to becoming a globally distributed exotic species in recent decades. It is generally accepted that island ecosystems offer weak biotic resistance to exotic species as compared to their native range. Here, we examined this empirically by comparing the relative competitive difference of *W. auropunctata* and locally dominant ants, between its native range of Mexico and introduced range of Puerto Rico. Resource competition was assessed between *W. auropunctata* and native ants under field conditions and in the laboratory. Furthermore, we compared resource competition at different temporal intervals ranging from short-term (< 2 hours) to long-term (14-days) dynamics. Our results are in contrast to common invasion predictions on island communities because we show that native species were resistant to *W. auropunctata* in its introduced range of Puerto Rico. We observed that the ground-foraging ant *Solenopsis invicta* competitive displaced *W. auropunctata* in Puerto Rico during short-term experiments. Meanwhile, the native arboreal ant *Linepithema iniquum* withstood competitive pressure from *W. auropunctata.* In the native range of Mexico, *W. auropunctata* was superior against *Solenopsis Picea* and *Pheidole protensa* species, but was outcompeted by dominant ants *Solenopsis geminata* and *Pheidole synanthropica.* This study challenges the relative importance of competitive ability in predicting invasion success. This is one of the few detailed comparative studies that examines exotic species performance between native and introduced habitats.

## Introduction

Biological invasions are important drivers of global change and pose a major challenge to biodiversity and human welfare (Vitousek 1990; Simberloff 2013). Determining how exotic species differ from native species is central to the study of invasion biology (Fridley *et al.* 2007; Lemoine, Burkepile & Parker 2016). Several factors including superior competitive abilities, natural enemy release, and anthropogenic disturbance are some of the major drivers behind species invasions (Lowry *et al.* 2013). A prevailing viewpoint, however, is that the success of invasive species is attributed to their superior competitive abilities relative to native species (Bruno *et al.* 2005). Invasions of exotic species provide a natural experiment for evaluating the relative roles of species interactions in driving community structure. In particular, understanding why some exotic species excel at dominating recipient communities requires a comparative approach between the native and introduced ranges of exotic species (Hierro, Maron & Callaway 2005).

In plant communities, competition by resident communities is a mechanism for biotic resistance to invasion (Mitchell *et al.* 2006). Within its native range, greater resident species richness and composition inhibits invasive species performance due to increased competition (Levine, Adler & Yelenik 2004). While positive correlations between native and invasive plant species richness have been reported at larger spatial scales (Stohlgren, Barnett & Kartesz 2003), this pattern appears to be correlated to spatial heterogeneities in abiotic conditions (Davies *et al.* 2005). In animal communities, interspecific competition is one of the mechanisms influencing ecological communities (Elton 1958; Schoener 1983). For instance, species-rich communities are thought to repel invaders from colonizing as residents use up all available resources and habitat (Case 1990; Drake 1991). It’s therefore important to test whether superior competitive abilities of invasive species explain their dominance in ecological communities.

Among animals, invasive ants are a prominent group to examine whether superior competitive abilities lead to the domination of native communities (Wittman 2014). In general, ant communities are thought to be structured by competition with behaviorally dominant species occupying large colony sizes with the ability to control access to limiting resources (Savolainen & Vepsäläinen 1988; Andersen & Patel 1994; Bertelsmeier *et al.* 2015). Globally, invasive ants are important and have been reported to negatively impact native ant communities, with five ant species considered to be among the worlds’ 100 worst invasive species (Lowe, Browne & Boudjelas 2000).

The little fire ant *W. auropunctata* is among the five worst invasive ant species (Wetterer & Porter 2003). In its introduced range, *W. auropunctata* has been found to negatively impact arthropods and vertebrate species, causing large reductions in their populations densities (Clark *et al.* 1982; Lubin 1984; Le Breton, Chazeau & Jourdan 2003; Vonshak *et al.* 2010; Wetterer 2013). In tropical invaded regions, high abundance levels of *W. auropunctata* have been associated with lower native ant diversity (Armbrecht & Ulloa-Chacón 2003). However, it’s unclear whether *W. auropunctata*’s superior competitive abilities are reflected at the community level. Furthermore, we lack comparative studies examining competitive dynamics of invasive species between their native and introduced ranges under natural conditions (Bertelsmeier *et al.* 2015). High abundance levels of *W. auropunctata*’s populations are found in human-disturbed habitats within its native and introduced ranges (Orivel *et al.* 2009a; Foucaud *et al.* 2009, 2010). *W. auropunctata* represents an excellent model system to tests whether competition serves as the principle mechanism for its dominance in human disturbed habits. We compared *W. auropunctata*’s competitive abilities between its native range of Mexico and introduced range of Puerto Rico. We tested whether *W. auropunctata*’s invasion success could be attributed to its competitive abilities in recipient communities. Comparing the relative rank of exotic species in local assemblages will allow us to make better predictions about global invasion patterns.

## Materials and Methods

### Study Sites

This study was conducted on an organic shaded agricultural coffee ecosystems within the native range of *W. auropunctata* populations in Mexico and in the introduced region of Puerto Rico. Both regions experience annual wet and dry seasons, as is common in many tropical regions. Data were collected during the wet season (summer) between the months of June and July in 2012, 2013, and 2014. In the Mexico site, a 45-ha plot was surveyed to map the spatial distribution of *W. auropunctata* colonies in a medium shaded organic coffee farm in the state of Chiapas in southern Mexico (15.1735835, -92.3382748), a 30-h a plot was surveyed in a low-shade conventional coffee farm (15.172465, -92.3301377), and finally a 6-ha plot was surveyed in a rustic part of the coffee farm with relatively high shade levels. We observed a total of 4 colonies in the 45-ha plot and 1 colony in the 30-ha plot ranging from 0.25-1 ha in size (Fig 1a). In Puerto Rico we surveyed 10 small coffee farms (mostly < 5 ha) in the mountainous regions of Puerto Rico within the municipalities of Orocovis, Lares, Adjuntas, and Utuado (18.175850 -66.4155700). The largest coffee farm (5-ha) where *W. auropunctata* populations occurred was located in Orocovis and this farm had the highest shade level (Fig 1b). For all ant surveys, species were identified in the field except in cases where species could not be readily identified. Un-identified individuals were later brought to the lab and identified to species or morpho-species.

**Fig 1.**
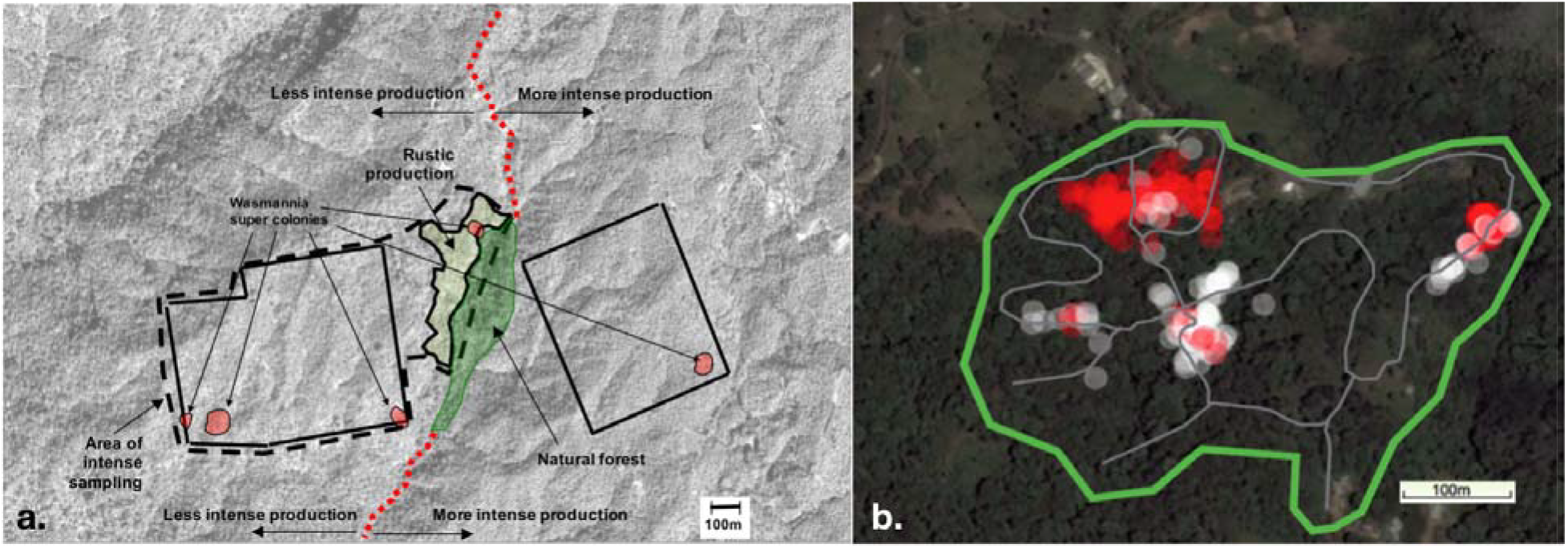
a). Survey of an intensively sampled area across approximately 80 ha of coffee farms. Shows the distribution of the five clusters of *W. auropunctata.* Bold lines indicate area already intensively sampled. Note that extensive sampling reveals the complete absence of *W. auropunctata* in all the area within the perimeter of the dotted lines, except for the five clumps indicated b). Spatial survey of *W. auropunctata* and *L. iniquum* distribution on a 6-ha Puerto Rican shaded coffee farm. A single megacolony of *W. auropunctata* is shown in the large clumped red region (~1ha). White and grey circles indicate smaller patches of *L. iniquum* interspersed with *W. auropunctata* nests (red). Thick gray lines were intensive sampled transects with baits placed every 2 meters.

### Natural History of the Study System

For our biogeographical comparison between *W. auropunctata* and resident ant species, we selected dominant ant competitors that naturally occurred across our coffee ecosystems and were found to nest in either the soil or in coffee plants. Within the native range of *W. auropunctata,* the most dominant ant species occupying sites in Mexico were *Solenopsis geminata, Pheidole protensa, Pheidole synantropica, and Solenopsis picea.* In the introduced range of Puerto Rico, the most dominant ant species we encountered were *Solenopsis invicta* and *Linipethema iniquum* (Fig 3).

**Fig 2.**
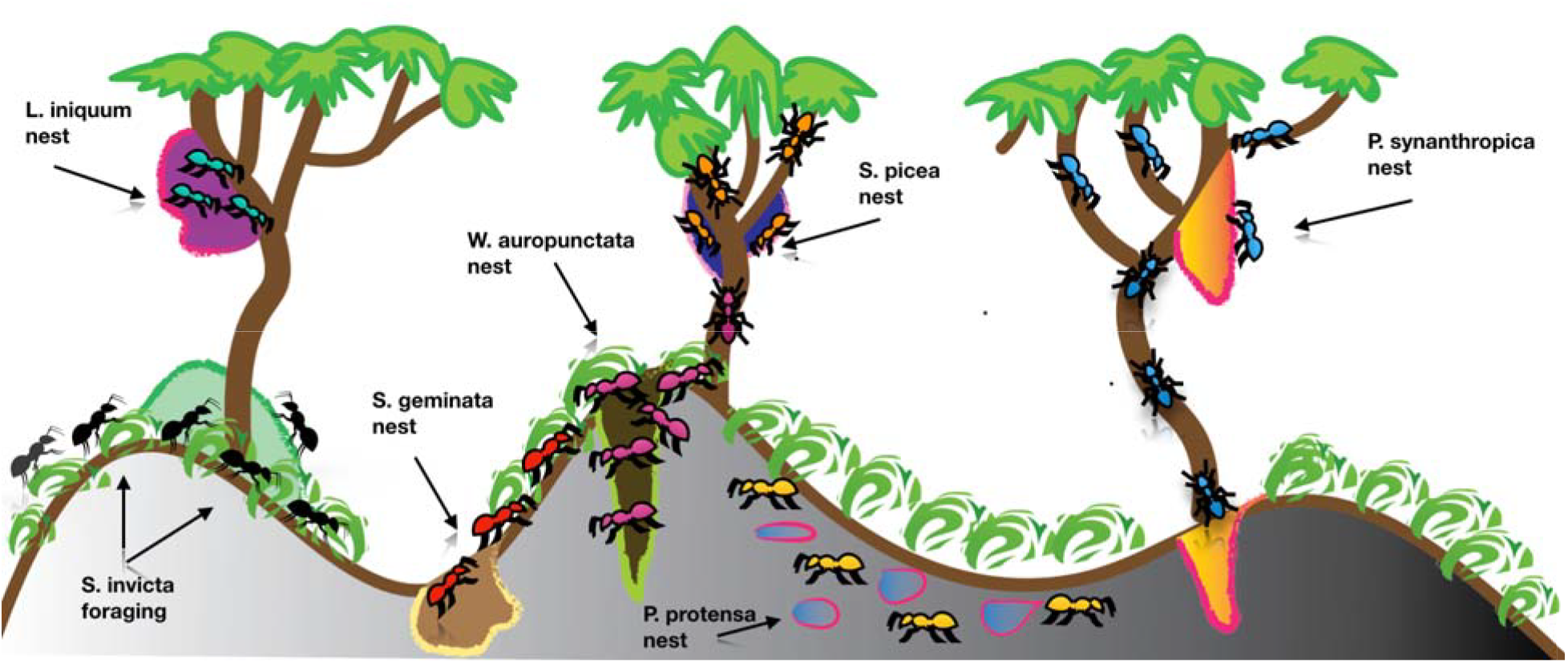
Representation of community structure of dominant ant species in Mexico and Puerto Rico. In Mexico, the soil nesting ants *S. geminata* (red) and *P. protensa* (yellow) are the dominant ground-foraging ants, while the arboreal ants *S. picea* (orange) and *P. synanthropica* (cyan) dominate in the trees. In Puerto Rico, the ground-foraging ant *S. invicta* (black) and the arboreal ant *L. iniquum* (blue) are dominant species.

**Figure 3.**
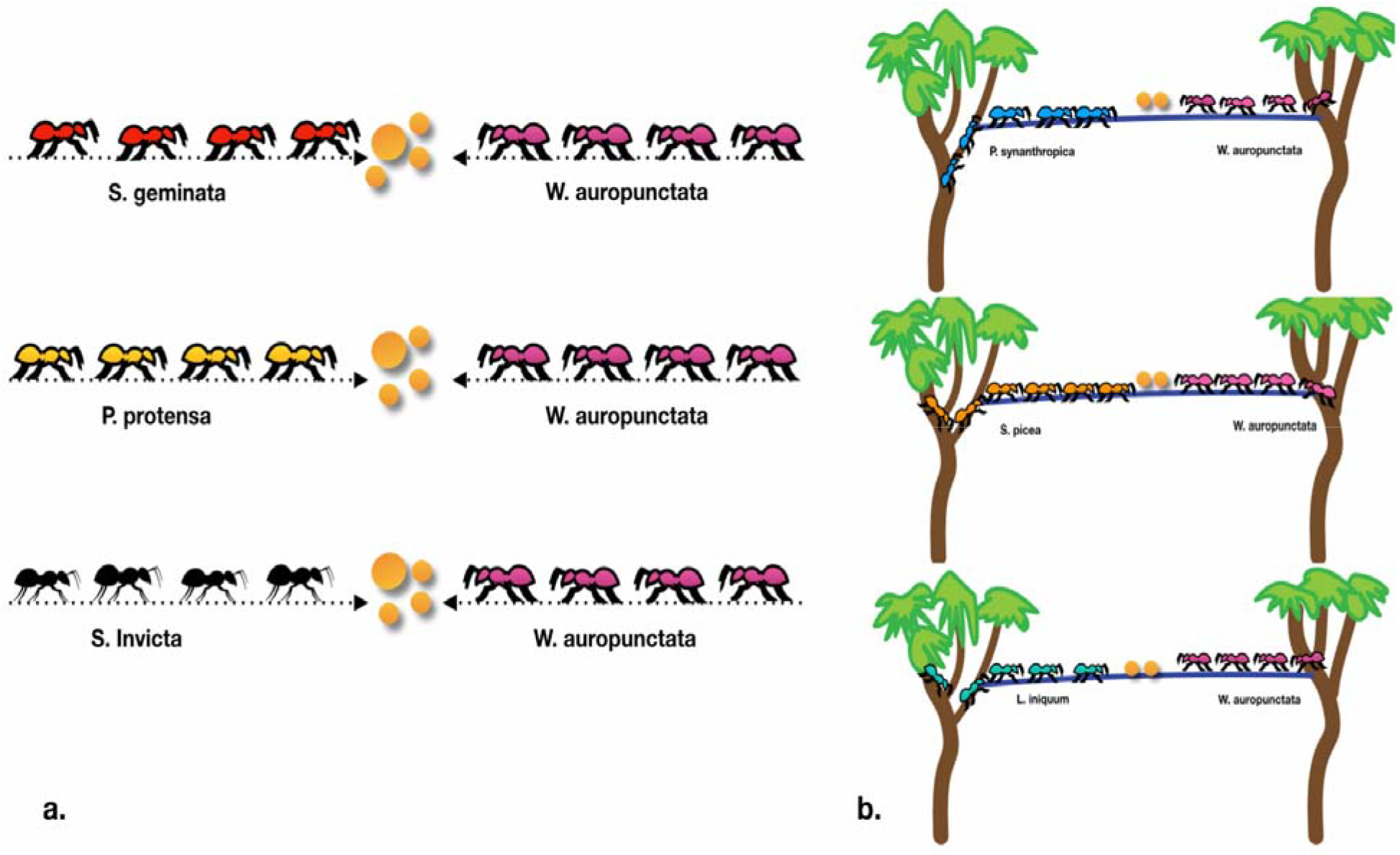
a) Field competition experiments on the ground were conducted by placing 10 baits in a 1-m line at the border between ant colonies and checked at 30-sec intervals for up to 80 minutes. Each of the 10 sites were separated by 10 cm within sites where species co-occurred. B) For field competition experiments on trees, we placed five baits on bamboo branches between coffee trees inhabiting ant nests. Each of the five sites was separated by 10 cm located between arboreal nesting sites.

The tropical fire ant *S. geminata* is one of the dominant ant species that occurs in our Mexico study site. *S. geminata* nests in the ground and is known for its aggressive behavior at baits where it uses it powerful sting to displace its competitors (Wetterer 2011). Its nests are characterized by a dome-shaped central chamber with covered tunnels that can withstand attacks from phorid fly parasites (Feener and Brown 2002, Morrison 2000). *S. geminata* commonly occurs in sunny areas as it can withstand higher temperatures (Torres 1984). Despite its ecological dominance, long-term competition experiments revealed that *S. geminata* was poor at discovering resources against *Pheidole* species (Perfecto & Vandermeer 2013).

*P. protensa* is one of the smallest and slowest moving ground-nesting ant on the farm. It forms many small nests with multiple entrances holes some of which belong to the same colony, while others are distinct from one another. *P. protensa* is a dominant ground-foraging ant that competes strongly for nest sites through scramble and contest competition (Vandermeer et al. 2010). In coffee, *P. protensa* serves as an important biological control agent against the coffee berry borer (Perfecto and Vandermeer 2013).

*P. synantropica* is a dominant ground-nesting ant that forages both on the ground and in coffee plants where it tends scale insect populations (Gonthier et al 2013). *P. synanthropica* has a large foraging capacity and is widely distributed throughout the coffee farm where it forages in large swarms and intensely competes with other ant species at baits (Ennis & Philpott 2017).

*S. picea* is a medium-sized black/dark brown arboreal ant that commonly nests under the bark of coffee trees. Where it occurs on the farm, *S. picea* can also be found nesting with other ant species such as *Crematogaster.* Its nests consist of polydomous and polygonous colonies which make it challenging to distinguish colony range in the field (antwiki.org).

In Puerto Rico, *S. invicta* is the most dominant ground-foraging ant found at our study site. *S. invicta* has in recent decades emerged as a global invasive species, including on the island of Puerto Rico where it is a behavioral dominant species with large colony sizes that competitively displace native ant species (Van Buren 1982). Its tolerance to human disturbed habitats combined with its painful sting makes this a notorious ant species (Wetterer 2013).

The arboreal ant *L. iniquum* is native to the eastern part of the Carribean region, including Puerto Rico, and is commonly found nesting in hollow twigs of coffee plants (Wheerler 1908, Wild 2008). *L. iniquum* appears to be polydomous, with many excavated nests not having queens at our study site. *L. iniquum* workers appear to be rapid at discovering resources in the field, but are also prone to phorid fly attacks (Yitbarek, personal observation).

### Ant surveys

Surveys were conducted to map the spatial distributions and abundance of dominant ant species. We walked dense trail systems by placing canned tuna baits on coffee plants at eye level (1 bait per coffee plant) approximately every 4 meters. Baits were checked every 30 minutes for the presence of ant species. A total of 1181 baits were placed within the 45-ha plot, 992 baits within the low-shade coffee farm, and another 229 baits within the rustic 6-ha plot (Fig 1a). Furthermore, we performed detailed ant surveys within 50 m x 50 m plots within the largest *W. auropunctata* dominated area on the farm to select competing dominant ant species for our experiments. (supplementary material A1). Detailed descriptions of the 45-ha coffee farm in Mexico have been reported elsewhere (Vandermeer et al. 2010).

In Puerto Rico, we placed a total of 664 baits on coffee plants across all 10 coffee farms (32 baits per ha, approximately 20 ha in total farm size). We did not sample ants on the ground because the majority of farms were located alongside steep hills that made sampling on the ground cumbersome (supplementary material A3). However, we did conduct detailed ground surveys in 20 m x 20 m plots within the largest farm in our study site in Puerto Rico from which we selected dominant ants from our competition experiments (supplementary material A2). Site locations in both Mexico and Puerto Rico were selected so that native ant species, both arboreal and ground foraging ants, overlapped in their ranges with *W. auropunctata* populations. Our baiting method did not set out to sample the entire community, but instead focused on dominant species that engage in competitive interactions (Armbrecht and Ulloa-Chacón 2003). Differences in farm size reflected the fact that coffee farms in the Soconoscu regions of Mexico are relatively large in size (300-ha) as compared to the central region of Puerto Rico, where the average farm size is considerably small (2-5 ha).

### Short-term competition experiment

We conducted short-term competition experiments between *W. auropunctata* and native ant species. In Mexico, competition experiments were performed on the ground between *W. auropunctata* and native ants *Solenopsis geminata* and *Pheidole protensa.* Arboreal competition experiments in turn were conducted with *Solenopsis picea* and *Pheidole synanthropica* species. In Puerto Rico, we conducted short-term experiments between *W. auropunctata* and the native ground-foraging ant *Solenopsis invicta.* In the lab, we conducted short-term competition experiments between *W. auropunctata* and the arboreal ant *Linepithema* iniquum.

For our terrestrial competition experiments, we recorded the timing of resource discovery, recruitment, and the total number of individual workers at baits by placing ten baits (i.e. tuna) in a 1-m line right at the border between ant colonies and subsequently checked the baits at 30-sec intervals for up to 80 minutes(Perfecto 1994). Each of the 10 sites was separated by 10 cm within sites where species co-occurred (Fig 3a). The intent of this experimental setup was to sample right at the border between mono-specific patches were competition took place. For our arboreal competition experiments in the field, we placed five baits on bamboo branches between coffee trees inhabiting ant nests. Each of the five sites was separated by 10 cm located between arboreal nesting sites (Fig 3b). We recorded the discovery time, recruitment time, and the total number of individual workers at baits.

### Long-term competition experiments

For the long-term competition experiments in Mexico, we used platforms to connect plastic containers inhabiting *W. auropunctata* and *S. picea* nests in a common foraging arena for 14-days. Holes were drilled in the plastic containers to allow ants from leaving the nest to the platform where the resources were placed. To prevent ants from escaping we placed the containers in a water bath. Prior to our laboratory experiment, we conducted field trials to identify potential dominant arboreal ants in the field. We selected *S. picea* nests due to their dominance in coffee plants and relative ease of nest collection. Caution was taking to collect nests that had sufficient amount of queens, workers, and brood in order to keep the nests viable in the lab. Each day the ants were supplied with sugar water and tuna in the lab to enable both species to compete for resources. We used 6 replicates in the experiment with different pairs of unique nest that had not previously been exposed to each other. Controls consisted of nests pairs that were not connected (n =2 pairs) with a platform. In Puerto Rico, we followed the same protocol for assessing the long-term competitive dynamics between *W. auropunctata* and the arboreal ant *L. iniquum.* We used 8 connected pairs of *W. auropunctata* and *L. iniquum* nests in the experiment and 2 pairs that served as control replicates.

### Analysis

For all the competition experiments, we analyzed the differences among species with respect to total number of workers at baits. Prior to conducting statistical analyses, we plotted the data to test for normality using the Shapiro-Wilk test. Because of the non-normality displayed in the data, we used the non-parametric Mann-Witney U-test with the native stats package in R (Team 2017). Nemenyi’s post-hoc tests were applied for pairwise multiple comparisons using the PMCMR package in R, v. 2.15. ANOVA analysis was performed to detect differences in the number of live ant workers found between treatments in the long-term competition experiments.

## Results

Within the native range, we found a low overall abundance of *W. auropunctata* surveyed in Mexico relative to its introduced range of Puerto Rico, where *W. auropunctata* was found to occur in much higher abundances. In the 45-hectare plot in Mexico (medium-shade organic coffee), we documented a total of 84 species, 3.4 % of which consisted of *W. auropunctata* found at baits. In the 30-hectare plot (low-shade conventional farm), we documented a total of 58 species of which only 0.4% occupied baits by *W. auropunctata.* In the 6-hectare plot (rustic coffee, within high shaded coffee), we detected *W. auropunctata* at 7.9 and 3.9% respectively at arboreal baits, whereas ground-baits occupancy by *W. auropunctata* was only 0.5%. While *W. auropunctata* distribution on the farm was limited in the native range, we detected a large cluster of *W. auropunctata* colonies that presumably make up part of a larger super-colony (Fig 1a). In Puerto Rico, we detected a total of 16 species throughout our surveys on 10 smaller-sized coffee farms (ranging from 1-6 ha). On average, we found 41.7 % of arboreal trees occupied by *W. auropunctata.* On the largest coffee farm (6 ha), we found that 82.2 % of arboreal trees were occupied by *W. auropunctata* while the second most dominant species *L. iniquum* occupied 17.8 % of trees. In most cases, *W. auropunctata* was widely distributed across all farms reaching high densities in patches it dominated thereby excluding other ant species from occupying nearby trees.

Overall, we found that *W. auropunctata* was patchily distributed between Mexico and Puerto Rico. The main difference, however, is that patches dominated by *W. auropunctata* included a greater diversity of native ant species in Mexico. For example, native species diversity in the largest patch (1-ha) ranged anywhere from 30-50 native ant species, with only 40 % of baits occupied by *W. auropunctata* (Fig 1a). In Puerto Rico, farms had on average less than 20 native ant species present, with the largest patch only containing 2 dominant arboreal ant species (Fig 1b).

### Short-term competition: Mexico and Puerto Rico

Competition between *W. auropunctata* and native ant species revealed important differences during contests. Interactions with *S. geminata* showed that *W. auropunctata* was able to rapidly recruit workers to baits during the first 20 minutes of the experiment (Fig 4 a), after which *S. geminata* took over and dominated the majority of baits by the end of the experiment (W = 598.5, p < 0.0001). Competitive interactions between *W. auropunctata* and *P. protensa* revealed oscillatory dynamics (Fig 4b). *W. auropunctata* rapidly increased at baits within the first 25-minutes after which *P. protensa* dominated for some time until getting surpassed by *W. auropunctata* (W = 1352., p < 0.0001). Arboreal competitive interactions between *W. auropunctata* and *S. picea* were equally intense. *S. picea* initially dominated the baits but was overtaken by *W. auropunctata* after 30-minutes (Fig 4 c). Despite *S. picea* maintaining a constant recruitment rate throughout the experiment, *W. auropunctata* steadily increased recruitment of workers enabling it to dominate most baits (W = 4190, p=0.0017). However, the arboreal ant *P. synanthropica* was competitively superior against *W. auropunctata* (Fig 4 d) during the experiment resulting in the complete dominance of baits (W = 1255.5, p < 0.0002). In Puerto Rico, we compared competitive dynamics with *W. auropunctata* and the dominant ant competitors *S. invicta* and *L. iniquum. W. auropunctata* initially increased its recruitment rate in response to *S. invicta* but was quickly overtaken by *S. invicta* as more of its workers dominated baits during the 50-min experiment. We didn’t detect a significant interaction in the competition experiment (W = 302.4, p=0.063), but we did observe that *S. invicta* was faster at discovering baits due to its rapid recruitment of workers (Fig 5a). In the laboratory, we compared short-term competitive interactions between *W. auropunctata* and *L. iniquum* (Fig 5b). *W. auropunctata* and *L. iniquum* populations oscillated throughout the experiment but the outcome of competition was indetermined (W = 4100, p=0.4724).

**Figure 4.**
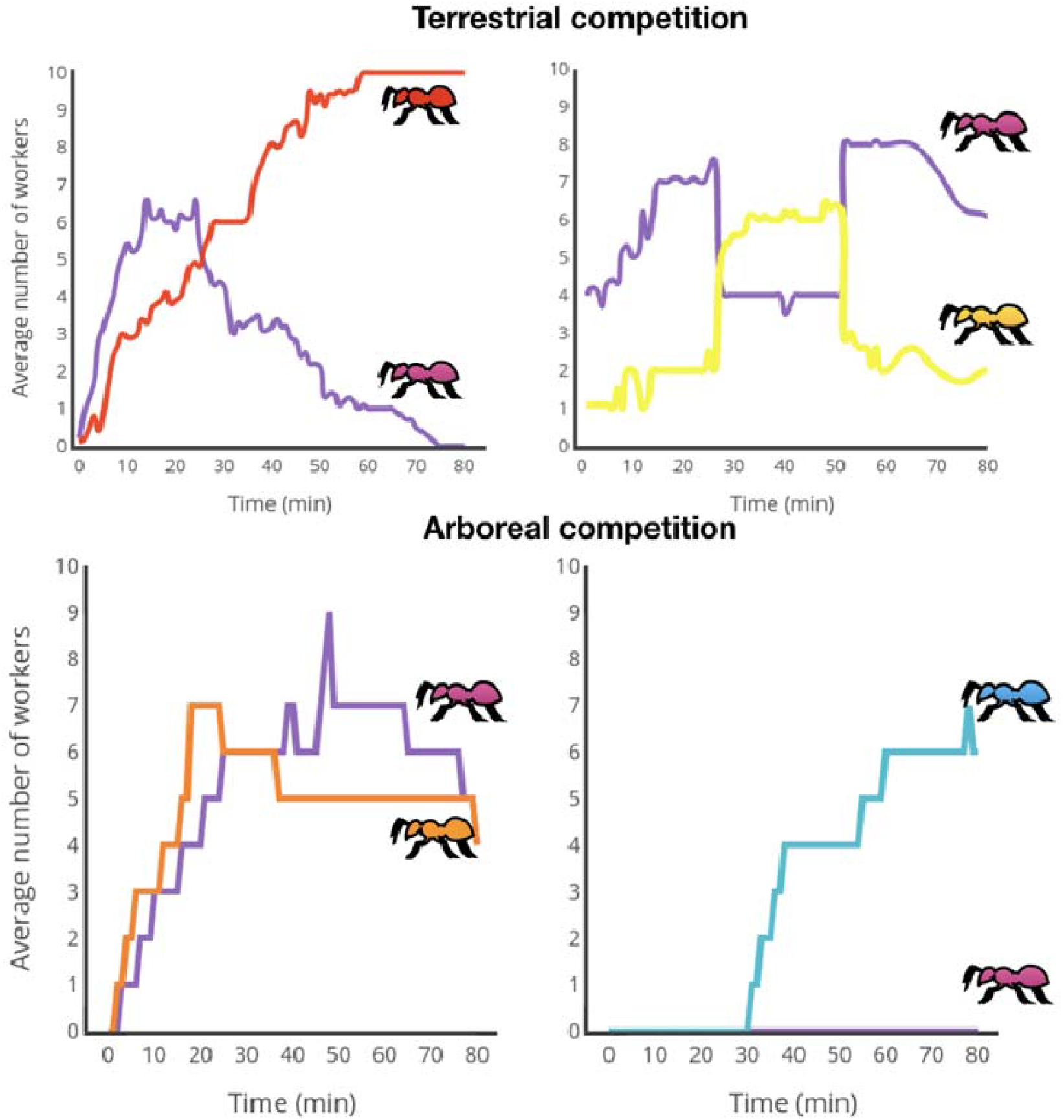
Average number of workers competing for resources between *W. auropunctata* (red) and native ant species in Mexico. Baits were placed at the boundaries of territorial patches (10 replicates). Top graphs interactions between *W. auropunctata* (purple) and ground foraging ants *S. geminata* (red) and *P. protensa* (yellow). *S. geminata* is a superior competitor (W = 598.5, p < 0.0001), but *W. auropunctata* displaces *P. protensa* on the ground (, p < 0.0001). Bottom graphs show interactions with the arboreal ants *S. picea* (orange) and *P. synanthropica* (cyan) (W = 4190, p=0.0017). *W. auropunctata* gains a competitive advantage against *S. picea,* but losses against *P. synanthropica* in the trees (, p < 0.0002).

**Figure 5.**
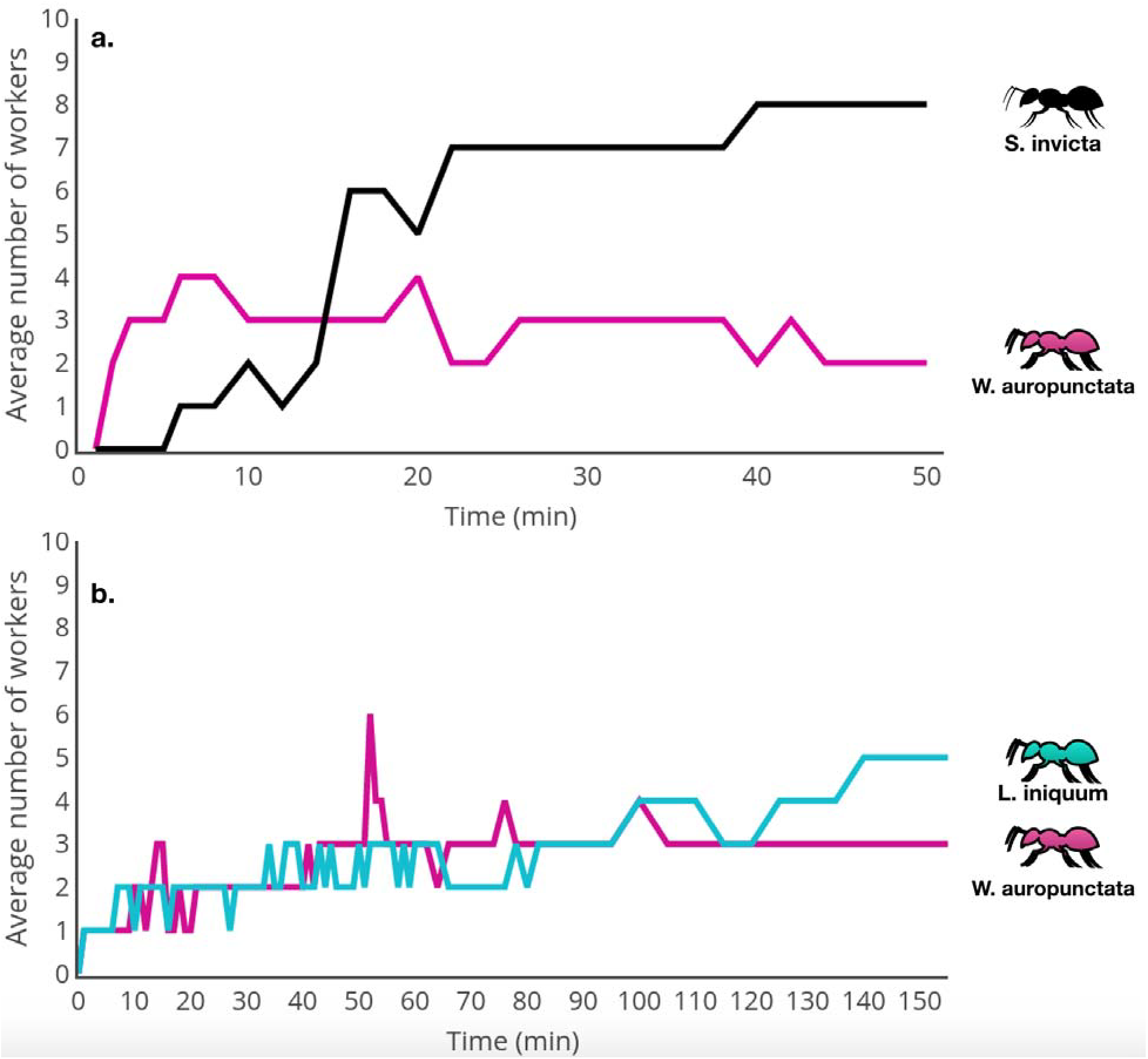
a) Field competition experiments conducted in Puerto Rico between *S. invicta* (black) and *W. auropunctata* (purple). By the end of the 50-min experiment, *S. invicta* was not competitively superior to *W. auropunctata* at baits (W = 302.4, p=0.063). b) Laboratory competition experiments between *L. iniquum* (blue) and *W. auropunctata* (purple). Both *L. iniquum* and *W. auropunctata* oscillated in their competitive interactions, with the final outcome of competition being undetermined (W = 4100, p=0.4724).

### Long-term competition: Mexico and Puerto Rico

*W. auropunctata* workers were superior against *S. picea* during the long-term experiment in Mexico (Fig 6a). We observed greater survival of *W. auropunctata* workers in the 14-day experiment (F=5.43, p=0.013). During the initial phase of the experiment, *S. picea* recruited higher number of workers to the foraging arena but preempted *W. auropunctata* workers from accessing resources. In most of the replicates, we observed that *W. auropunctata* had a clear numerical advantage over *S. picea.* In five of our replicates, *W. auropunctata* invaded the nest boxes of *S. picea* resulting in a high worker density of *W. auropunctata* including many brood and queens. In one replicate (#3), we observed a nest invasion by *W. auropunctata* workers into a *S. picea* nest, with queens and brood moved into the invaded nest. Further inspection into nests with a tiny camera revealed that *W. auropunctata* workers frequently dispersed their brood and queens within nests. In Puerto Rico, the number of live *W. auropunctata* workers was not significantly higher than *L. iniquum* (F=1.87, p=0.17). In the first two days, *L. iniquum* was much faster at recruiting workers to baits compared to *W. auropunctata.* In about one quarter of the replicates, few *L. iniquum* workers were found alive in the nest boxes as they were killed by *W. auropunctata* workers. We did observe occasional fighting between species, but *L. iniquum* was quick enough to avoid capture by *W. auropunctata* workers in the foraging arena (Fig 6b).

**Figure 6.**
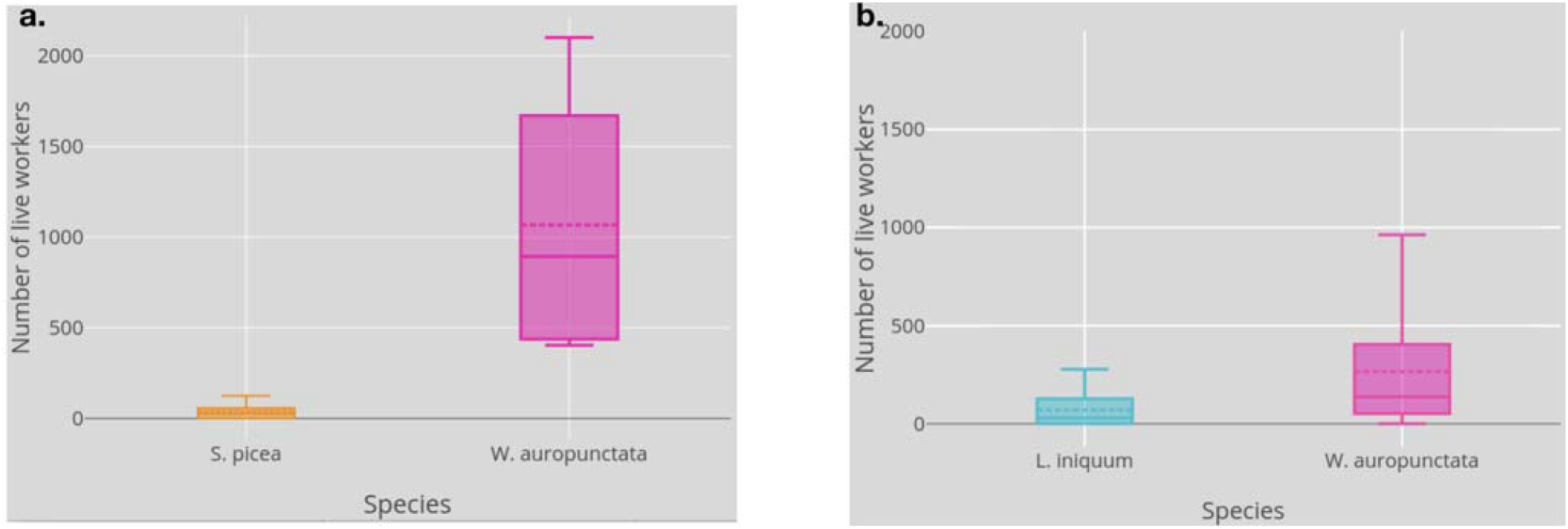
a) Long-term competitive interactions in Mexico between ground-foraging ants *W. auropunctata* (purple) and *S. picea* (orange). The average number of live ant workers recorded at the end of long-term experiment (14-days). *W. auropunctata* has a higher number of workers found in nest boxes, including those workers that invaded *S. picea* nests (F=5.437, P= 0.013). b) Long-term laboratory competition experiment between arboreal ants *W. auropunctata* (purple) and *L. iniquum* (cyan) in Puerto Rico. The average number of live ant workers recorded at the end of the long-term experiment (14-days). We did not detect significance differences between live *W. auropunctata* and *L. iniquum* workers (F=1.87, P=0.17) Error bars are standard errors of mean.

## Discussion

One of our main findings is that locally diverse ant assemblages resisted *W. auropunctata* in both Mexico and Puerto Rico. We identified several dominant species from the local ant assemblage in the native and introduced habitats and compared their competitive interactions with *W. auropunctata.* In particular, we examined resource and interference competition across short and long-term time scales. We found that in Mexico the ground-foraging ant *S. geminata* and the arboreal ant *P. synanthropica* were superior against *W. auropunctata.* In contrast, the ground-foraging ant *P. protensa* and the arboreal ant *S. picea* were outcompeted by *W. auropunctata.*

Furthermore, laboratory experiments showed that *W. auropunctata* was superior against *S. picea* during long-term competition experiments. In Puerto Rico, another exotic ant *S. invicta* outcompeted *W. auropunctata* in areas where they overlapped. Laboratory experiments with the native arboreal ant *L. iniquum* showed that *W. auropunctata* was not superior at baits across the short and long-term competition experiments.

Exotic ants can displace native ant species through interference and resource competition (Sakai et al. 2001). While these traits have been implicated in the success of exotic ants, they don’t always explain why exotic ants excel at displacing native ant species (Holway et al. 2002). Within native communities, competitive trade-offs between behavioral dominance and resource discovery are thought to promote species coexistence (Fellers 1987; Adler et al. 2007; Bertelsmeier et al. 2015). Exotic ants can violate the discovery-dominance trade-off in the introduced range, in the case of Argentine ants, resulting in a competitive advantage over native species (Holway 1998). While the breaking of the discovery-dominance tradeoff has been linked to the invasive success of Argentine ants, we lack such information on other major invasive ant species. Laboratory experiments involving *W. auropunctata* and native ant species, showed that *W. auropunctata* had the poorest foraging abilities in terms of resources discovery (Vonshak et al. 2012). In previous work, we compared competitive trade-offs and found that *W. auropunctata* was less efficient in discovering and recruiting worker to baits in their introduced habitat of Puerto Rico relative to their native habitat in Mexico, suggesting that competitive trade-offs are not indicative of invasive success (Yitbarek, Vandermeer & Perfecto 2017).

The organization of exotic species within local species assemblages in their native ranges allows us to understand why some exotic species thrive in their introduced ranges (Calcaterra et al. 2016). Here we sought to understand how *W. auropunctata* ranks relatively to locally dominant native ant species. In Mexico, several native ant species from the local species assemblage were able to withstand *W. auropunctata.* The tropical ant *S. geminata* showed high levels of aggressiveness towards *W. auropunctata* at baits. Although *W. auropunctata* was initially able to dominate resources, *S. geminata’s* high recruitment and interference ability led to the displacement of *W. auropunctata.* Although *S. geminata* is frequent victim to attacks by phorid flies which can interfere with its ability to rapidly discover resources(Brown and Feener 1991; Morrison 2000), we didn’t detect phorid flies at our site which greatly enhanced *S. geminata’s* dominance at baits. In comparison, *P. protensa* is a slow moving ground forager ant with many small nests that are widely distributed. Although *W. auropunctata* was initially dominant at baits, *P. protensa* was able to overtake the baits for some time after which *W. auropunctata* regained its dominance again. These cyclical patterns are illustrative of the transient nature of competitive interactions.

The arboreal ant *S. picea* is a minute ant that primarily forages on coffee where its nests can be found under the bark on the trunk of the coffee tree. *S. picea* was initially more dominant but as *W. auropunctata* recruited its workers from the leaf litter and arboreal nests to the baits it started to regain its dominance. The ability to forage and nest on the ground and the trees enables *W. auropunctata* to hold a competitive advantage over native species. We confirmed this pattern in the long-term experiment were *W. auropunctata* interference ability enabled it to invade S. picea nests. *P. synanthropica* species nest in the ground but forage both in the leaf litter and in the coffee trees. *P. synanthropica* was competitively dominant over *W. auropunctata* as it quickly discovered baits and vociferously defended against *W. auropunctata* intruders. *P. synanthropica’s* dominance has also been observed in competition experiments against *S. geminata* and *P. protensa* (Perfecto and Vandermeer 2013).

In Puerto Rico, the exotic ant *S. invicta* was superior to *W. auropunctata* at ground baits. *S. invicta* reached high population densities at baits which forced *W. auropuncatata* to being pushed out at baits. While *S. invicta* attained a numerical advantage over *W. auropunctata,* the extent of their colonies was limited to areas of the farm that received more sun, which allowed *W. auropunctata* to retain its dominance in other areas where *S. invicta* did not occur. Laboratory experiments showed an oscillatory dynamics between *W. auropunctata* and *L. iniquum* competition. We found that *L. iniquum* workers were faster at discovering resources but were less efficient at guarding baits against *W. auropunctata* workers.

Although we found a higher number of live *W. auropunctata* workers during the longterm experiment, the patterns were not significantly different from *L. iniquum* workers. *W. auropunctata* displaced *L. iniquum* by raiding their nest and killing its workers and brood, in some cases moving its entire nest into the invaded species. However, *L. iniquum’s* rapid tempo made it challenging for slowing moving *W. auropunctata* to find them. Our long-term experiments show that nest invasions are a potential mechanism by which *W. auropunctata* displaces native ant species. Previous laboratory experiments showed that *W. auropunctata* displaced the native ant species *Monomorium subupacum* by invading its nest and consuming workers and brood (Vonshak et al. 2012). Similar evidence for nest raiding of ant colonies has been found for the invasive ants *L. humile* and *P. megacephela* in their introduced ranges (Holway et al. 2002; Zee and Holway 2006; Dejean et al. 2008).

Recent evolutionary studies have shown that *W. auropunctata* invasion displays a polymorphism in its reproductive system (Foucaud et al. 2010; Rey et al. 2013). This shift in reproductive structure first occurred within the native range of *W. auropunctata* from low-density populations in natural habitats (i.e. sexual populations) to high-density workers (i.e. clonal populations) in anthropogenic habitats (Fournier et al. 2005; Foucaud et al. 2009, 2010). This adaptation to human modified habitats suggests that *W. auropunctata’s* invasive success is associated to its clonal reproductive structure (Hufbauer et al. 2012). It remains unclear, however, to what extent clonality in the introduced range contributes to ecological dominance in *W. auropunctata* populations. Our study compared the invasion dynamics of *W. auropunctata* populations in agricultural ecosystems. *W. auropunctata* populations occurring in these anthropogenic sites are mostly likely clonal (Orivel et al. 2009a). Within our most densely populated site in Mexico, we found that *W. auropunctata* was competitively superior against several dominant ant species, contrary to Puerto Rico where the most dominant ant *L. iniquum* was able to withstand competitive pressure by *W. auropunctata.*

Although direct competitive interactions can structure local assemblages, indirect interactions can alter competitive outcomes (Hsieh and Perfecto 2012). Trait-mediated-indirect effects (TMII) have been found to play an important role in structuring ecological communities (Werner and Peacor 2003). Phorid parasitoids can be highly host specific, by using ant pheromones, which as a consequence reduce the number of ant foragers at resources (Feener 2000). This can have important ramifications in ecosystems when phorids reduce competitive hierarchies (Hsieh and Perfecto 2012). Several studies have reported that Pseudacteon phorids affect resource retrieval by *S. geminata* species (Feener and Brown 1992; Morrison 1999). In Mexico, we didn’t observe phorids during direct encounters involving *S. geminata* and *W. auropunctata* but that may be heightened by bio-geographical disparities (Feener et al. 2008). Similarly, reductions in foraging activities of *Linipethema* species have been recorded in the presence of phorid species (Orr et al. 2003). Future studies should evaluate whether *L. iniquum* species get attacked by phorid flies in the field and how this indirect effect impacts competitive interactions with *W. auropunctata* in Puerto Rico.

A central question in invasion biology is to what extent exotic species are limited by their competitors in their native ranges and how release from interspecific competition can explain their invasive success in the introduced range. We found that *W. auropunctata* was able to outcompete dominant ants in Mexico, but that it was restricted by native competitors in its introduced range of Puerto Rico. These findings suggest that release from interspecific competition does not explain the invasive success of *W. auropunctata* in novel environments. An alternative explanation for *W. auropunctata’s* invasive success is the escape from natural enemies in the introduced range (Torchin et al. 2003). Recent work suggests that a purge from endosymbiotic bacteria might have contributed to *W. auropunctata’s* invasion success because of lower parasite pressure in the introduced range (Rey et al. 2013). The loss of parasites during invasion has been reported for several exotic ant species (Reuter et al. 2004; Yang et al. 2010).

In the case of *W. auropunctata,* it would be interesting to compare the prevalence of endosymbiotic bacteria within *W. auropunctata* populations between Puerto and Mexico. In particular, experiments that seek to determine how changes in *W. auropunctata’s* microbiota influence interspecific interactions with native species.

